# Robust dual dynamics of ictogenesis in temporal lobe seizures in vivo

**DOI:** 10.1101/2025.11.02.686119

**Authors:** Amir Mashiel, Shai Achvat, Yitzhak Schiller

## Abstract

Ictogenesis is a pivotal yet poorly understood aspect of epilepsy. Using two-photon calcium imaging, we simultaneously recorded excitatory CA1 pyramidal neurons (PNs) and inhibitory parvalbumin and somatostatin interneurons (INs) to examine seizure initiation dynamics in acute and chronic temporal lobe epilepsy models in vivo. Acute seizures exhibited two initiation patterns: INs-first and simultaneous PNs-INs activation, while chronic spontaneous seizures followed only the latter. Chemogenetic and optogenetic experiments further supported dual initiation dynamics, with seizures arising from either PNs or INs activation. At the onset of seizures, synchronization of INs exceeded that of PNs. The recruitment rank order of PNs, and to a lesser extent INs, remained highly consistent across seizures, indicating a stereotyped, robust initiation pathway for seizure initiation. These findings indicate seizures can emerge via two distinct yet stable pathways, reflecting dynamic but stereotyped initiation patterns shaped by excitatory-inhibitory interactions within epileptic networks.

## Introduction

Epilepsy is a common neurological disorder affecting ∼1% of the population and is defined by recurrent seizures^1,2^. In epilepsy, the cortical network transitions between two states: the asymptomatic interictal state, occasionally marked by brief abnormal EEG discharges, and the symptomatic ictal state of seizures. The interictal-to-ictal state transition, termed ictogenesis, underlies seizure initiation and is the primary target of pharmacological and neurostimulation anti-seizure therapies^1^.

The mechanisms driving temporal lobe seizure initiation have been extensively studied in vitro using brain slices and whole-brain preparations. These studies often report that inhibitory interneuron (IN) activation precedes that of excitatory pyramidal neurons (PNs), suggesting that seizure onset may result from hyperactivation of INs^1,3,4^. This is supported by findings that optogenetic activation of INs can trigger seizures^5–11^. The precise mechanisms through which optogenetic IN activation leads to seizure initiation remain unclear and may be related to post-inhibition rebound activation of PNs^12,13^, or changes in the intra- and extracellular ionic concentrations including accumulation of extracellular potassium or intracellular chloride^3,4,14,15^.

Another key aspect of ictogenesis examined in vitro is the consistency of neuronal recruitment at seizure onset. A notable study using hippocampal slice cultures showed that seizure initiation is stochastic and non-stereotypic, with different neuronal recruitment patterns across seizures—suggesting seizures emerge from a pathological network rather than a fixed subset of neurons^16^.

Despite these insights, most ictogenesis studies have been conducted in vitro. In vivo cellular-resolution studies remain limited, and none have utilized two-photon calcium imaging, which allows precise cell-type identification and high spatial resolution. Additionally, prior work has primarily focused on acute chemoconvulsant-induced seizures, with little exploration of ictogenesis during spontaneous seizures in chronic epilepsy models. Notably, in vivo studies of neocortical seizures have failed to observe early IN activation at seizure onset, even though both neocortical and temporal lobe seizures typically begin with low-voltage fast EEG activity^17^.

In this study, we used two-photon calcium imaging to simultaneously monitor CA1 excitatory PNs and PV- or SST-expressing INs during seizure initiation in vivo. We addressed two key questions: (1) What are the dynamics of excitatory/inhibitory (E/I) interactions at seizure onset? and (2) How consistent is neuronal recruitment across seizures? To answer these, we employed two in vivo models of temporal lobe epilepsy: acute 4-aminopyridine (4-AP)-induced seizures and spontaneous seizures in the chronic kainic acid (KA) model.

## Materials and Methods

### Experimental Design

We performed two-photon calcium imaging in awake, head-restrained mice to study seizure initiation in the hippocampus during acute and spontaneous seizures. GCaMP6f was expressed in CA1 pyramidal neurons (PNs), either alone or with PV- or SST-expressing interneurons (INs). To allow imaging a chronic hippocampal window was constructed as previously described^18^.

### Mice and Surgical Procedures

All procedures adhered to NIH and Technion IACUC guidelines. Adult (8–12 week) C57BL/6 WT or transgenic mice (Thy1-GCaMP6f^19^, PV-Cre^20^, SST-Cre^21^ were used. Mice were kept on a 12:12 h reverse light-dark cycle. Surgery was performed under isoflurane, with ketoprofen and buprenorphine for analgesia.

A 0.5 mm craniotomy was made above left CA1 (2.2 mm posterior, 1.35 mm lateral to Bregma). AAVs expressing GCaMP6f^22^ (CamKII-driven for WT; mix with Flex-mRuby2-GCaMP6f for PV/SST-Cre) were injected (350 nL) at 1.35 mm depth using a hydraulic micromanipulator. A second 2.77 mm craniotomy allowed aspiration of cortex up to the hippocampal alveus. A cannula with a perforated 170 µm thick glass coverslip at the bottom was inserted, sealed, and secured with dental cement; a headpost was affixed.

### Two-Photon Calcium Imaging

Two photon imaging was performed as previously described^17,23,24^. We used an Ultima 2P plus microscope (Bruker) with a resonant scanner and 16X water immersion objective (NA 0.8, Nikon). Images were acquired at 30Hz, or 12–15Hz for optogenetic and 10–12Hz for SST recordings. Dual-plane imaging was enabled by an electrically tunable lens (ETL). Each trial lasted 60 s with 10 s inter-trial intervals; 25–40 trials were acquired per session.

### Seizure Induction

Acute seizures were induced by applying 4-AP (4.5 mM in aCSF) topically through the cannula window. In chronic models, KA (20–50 nL in saline) was injected at 1.8 mm posterior, 1.6 mm lateral, and 1.5 mm depth. Seizures initially developed, and then few weeks later spontaneous seizures developed in most mice. Only mice with ≥1 seizure/h were imaged.

### Chemogenetic and optogenetic experiments

To chemogenetically inhibit INs activity, AAV2-hSyn-DIO-hM4Di-mCherry was injected (150–180 nL) into CA1 of PV-or SST-Cre mice. Clozapine-N-oxide (CNO, 5 mg/kg, IP) was administered 20 minutes prior to imaging. After control trials, 4-AP was added to evoke seizures.

To optogenetically activate INs PV- or SST-Cre mice were injected with AAV1.CAGGS.Flex.ChR2-mRuby2. WPRE.SV40. To activates PNs wildtype mice were injected with AAV1.CAMK2.ChR2-mRuby2. WPRE.SV40 ChR2-expressing neurons were activated with 10 ms pulses of 470 nm light at 10–15Hz at 10 mW intensity.

### Electrophysiology

Local field potentials (LFPs) were recorded from CA1 using a stainless-steel electrode inserted contralaterally via a 0.6 mm craniotomy (AP 0 mm, ML 2.31 mm) and angled at 35° to reach below the imaging field. Signals were acquired at 10kHz (MultiClamp 700B), band-pass filtered (1–70Hz), and synchronized with imaging via TTL pulses.

### Histology

At the experiment’s end, mice were perfused with PBS followed by 4% paraformaldehyde, and later brains were sectioned at 100 µm, and imaged with a Pannoramic slide scanner (3D Histech).

### Quantification and analysis

The fluorescence data acquired were first registered to correct for brain motion artifacts as previously described^17,23,24^. Regions of interest (ROIs) were detected manually, and the pixels within each ROI were averaged for every frame. ΔF/F was computed as previously described^17,23,24^

Analysis of data was performed using homemade MATLAB (MathWorks, Natick, MA). Our analysis included averaging of the calcium transients; calculating the pairwise Pearson correlation coefficient between individual neuronal pairs and averaging the individual results across all neuronal pairs in individual mice; determining the onset time of activation of individual neurons; calculating the rank order of recruitment of the different neurons during individual seizures and the inter-seizure Spearman correlation coefficient of the recruitment rank order of the different seizures within individual mice.

Data collected in this project will be stored in the figshare.com database. The code of the homemade analysis programs will be stored in github.

## Results

### Calcium imaging of epileptic seizures in the CA1 region of the hippocampus

To explore the dynamics of excitatory CA1 PNs during seizure initiation we combined in vivo two-photon calcium imaging from CA1 PNs expressing the genetically encoded calcium indicator GCaMP6f with electrophysiological local field potential (LFP) recordings during seizures. GCaMP6 was expressed in excitatory CA1 PNs using two methods: AAV viral vectors expressing GCaMP6 under the control of the CaMK2 promoter^25^ and a transgenic mouse line expressing GCaMP6f in CA1 neurons^19^ (Fig. 1a-c). LFP was recorded using a metal electrode implanted in the stratum radiatum beneath our imaging field (fig. 1a).

**Figure 1.**
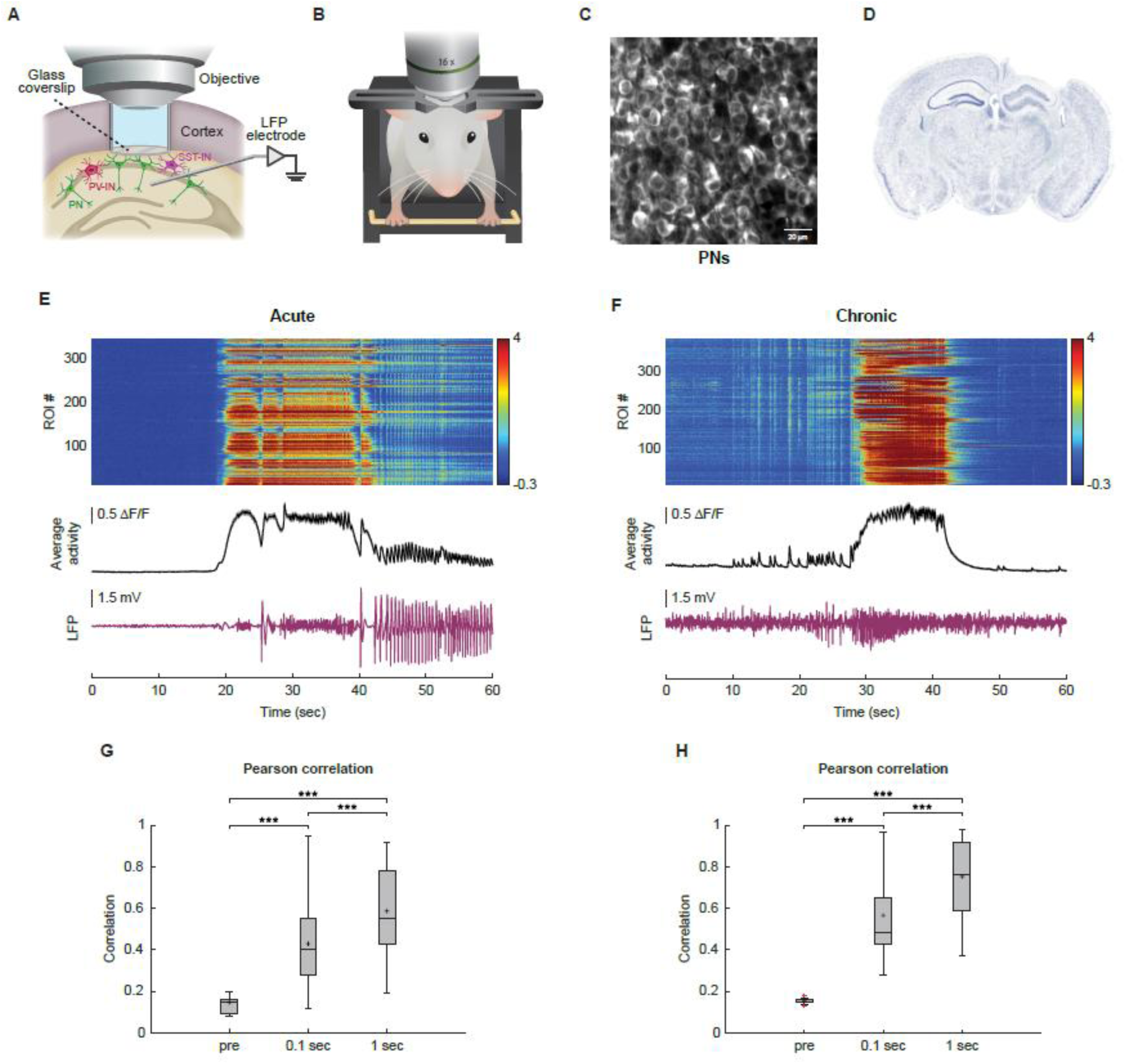
Activation of excitatory pyramidal neurons during seizures in acute and chronic models of epilepsy. **a,** Schematic illustration of the surgical procedure and imaging setup: a cannula is implanted over CA1 region and an LFP electrode placed in the CA1 region below the cannula. Excitatory PNs (green) and inhibitory INs are shown in the CA1 region below the imaging window. Imaging is performed through a water-immersed objective positioned above the cannula **b,** Schematic illustration of the imaging setup demonstrating a head-fixed mouse under two-photon microscopy. **c,** Two-photon imaging of GCaMP6f-expressing PNs in stratum pyramidale of CA1 in a mouse with acute 4-AP induced seizures. **d,** Coronal histological section of a mouse brain after KA injection to the left CA1. Note the damaged left hippocampus compared to the uninjected contralateral side. **e,** Combined recording of calcium imaging from CA1 PNs and LFP recording during an acute 4-AP induced seizure. The upper panel shows the calcium fluorescence transients presented as ΔF/F of the individual recorded PNs (over 300) during the seizure. The calcium transients were recorded using the calcium indicator GCamP6f and presented as color coded ΔF/F responses. Black middle trace shows the average (mean±SD) ΔF/F response over all recorded ROIs. Magenta lower shows the simultaneously recorded LFP response. **f,** Same as **e** for a spontaneous seizure recorded in the chronic KA model. **g,** Box plots of the average maximal Pearson correlation coefficient calculated for three time-windows in acute 4-AP induced seizures, immediately before seizure initiation (-1.2 sec up to -0.2 sec prior to seizure onset), during the initial 100 msec of seizures, and during the initial 1 sec of seizures. For these analyses Pearson correlation coefficients were calculated for each neuronal pair during the different time-windows, and then averaged over all neuronal pairs. **h,** Same as **g** for spontaneous seizures in the chronic KA model. PNs = pyramidal neurons, ROI = region of interest; ***p < 0.001; transverse lines mark the median values, black crosses mark the mean values, and red crosses mark outliers.

We investigated two different epilepsy models. Firstly, acute seizures induced by locally applying the potassium channel blocker 4-AP^26–28^. This model allowed for a controlled seizure onset site and robust seizure generation. Secondly, we explored the chronic KA model^29^ (Fig. 1d), where spontaneous seizures develop several weeks after local KA injection into the hippocampus. However, these spontaneous seizures are unpredictable and with limited knowledge of the seizure onset site within the hippocampus.

Our experiments demonstrated rapid, synchronized recruitment of CA1 PNs at seizure onset in both the acute 4-AP and chronic KA models (Fig. 1e-h). Moreover, the averaged calcium transients consistently marked the seizure onset time, as confirmed by the simultaneous LFP recordings (Fig. 1e,f).

### Relative activation of excitatory pyramidal neurons and inhibitory interneurons during onset of 4-AP induced seizures

While it is widely acknowledged that E/I imbalance plays a critical role in seizure initiation, prior studies have reported variable results concerning the relative recruitment of excitatory PNs and inhibitory INs during seizure onset, especially in the hippocampus^1,3,4^. To directly investigate this question, we conducted simultaneous recordings from excitatory PNs and inhibitory INs during 4-AP-induced seizures. We focused on two subtypes of INs: axo-somatic targeting PV-expressing INs, and dendritic targeting SST-expressing INs^30,31^. To simultaneously record from PNs and either PV- or SST-expressing INs, we utilized PV- or SST-Cre transgenic mice. We injected two different AAV viral vectors into their CA1 region: the first expressed GCaMP6f under the control of the CaMK2 promoter in PNs, and the second expressed flexed GCaMP6f and the red fluorescent protein mRuby2 under the control of the synapsin promoter, and thus expressed only in INs in a Cre-dependent manner (Fig. 2a-c). Consequently, GCaMP6f was expressed in both PNs and INs, while only INs co-expressed the red fluorescent protein mRuby2 as well (Fig. 2c). Given our previous experiments demonstrated the composite calcium transient reliably reports seizure onset (Fig. 1e,f), we refrained from simultaneously recording the LFP to minimize damage caused by intra-hippocampal metal electrodes.

**Figure 2.**
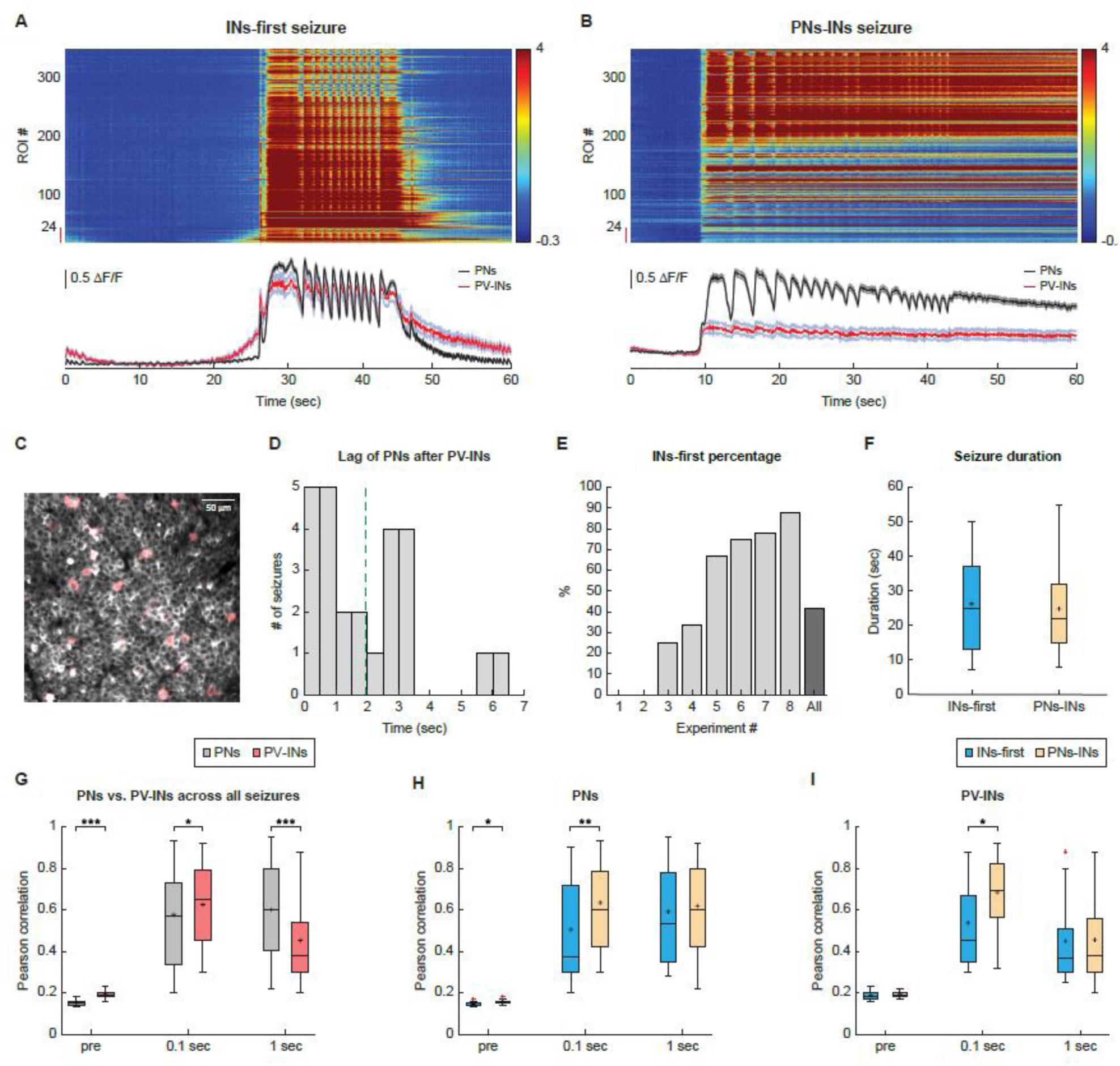
Simultaneous recordings of excitatory pyramidal neurons and inhibitory PV-expressing interneurons reveal dual dynamics during initiation of acute 4-AP induced seizures. **a,b,** The calcium fluorescence transients presented as ΔF/F for the individual PV-expressing INs (lower 24 ROIs marked by a red vertical line on the left) and PNs (all remaining ROIs) during two individual acute 4-AP induced seizures in the same mouse. The calcium transients were recorded using the calcium indicator GCamP6f and presented as color coded ΔF/F responses. The panels below show traces of the average (mean±SD) ΔF/F response over all recorded PV-expressing INs (red trace) and all simultaneously recorded PNs (black trace). Note that in the seizure shown in panel a, activation of PV-expressing INs preceded that of PNs (INs-first initiation dynamics), while in the seizure shown in panel b, activation of PNs and PV-expressing INs occurred simultaneously (simultaneous PNs-INs initiation dynamics). c, Overlaid two-photon images of GCaMP6f which was expressed in both PNs and PV-expressing INs (gray), and of mRuby2-postive PV-expressing INs (red). d, A histogram showing the latency between initial firing of PV-expressing INs and PNs across all seizures with INs-first initiation dynamics. Dashed line marks the median value. e, A histogram of the percentage of seizures with INs-first initiation dynamics in the 8 different recorded mice (light gray bars) and average value for all mice (dark gray right bar). f, Box plots of seizure duration of seizures with INs-first and simultaneous PNs-INs initiation dynamics. g, Box plots of the average Pearson correlation coefficients of PNs compared to PV-expressing INs during different time windows of seizures. h, Box plots comparing the average Pearson correlation coefficients of PNs during different time windows of seizures with INs-first (blue) and simultaneous PNs-INs (yellow) initiation dynamics. i, Same as h for PV-expressing INs. PNs = pyramidal neurons, PV = parvalbumin, INs = interneurons, ROI = region of interest; *p < 0.05, **p < 0.01, ***p < 0.001; transverse lines mark the median values, black crosses mark the mean values, and red crosses mark outliers.

Simultaneous recordings of CA1 PNs and PV-expressing INs during 4-AP induced seizures revealed two fundamentally different patterns of initiation dynamics. In 58.6% of seizures (36 seizures in 8 mice), PV-expressing INs and PNs were activated simultaneously within our temporal resolution limitations (simultaneous PNs-INs initiation dynamics). In the remaining 41.4% of seizures (25 seizures in 6 mice), activation of PV-expressing INs preceded activation of PNs by an average of 1.91±1.67 seconds (INs-first initiation dynamics) (Fig. 2a,b,d,e). Two out of eight mice (25%) showed only a single initiation dynamics pattern in all their seizures. Yet, in most mice (6 out of 8), seizures alternated between the two initiation dynamics patterns (Fig. 2e).

Temporal lobe seizures show two main EEG onset patterns, low voltage fast activity (LVF) and large amplitude hypersynchronous (HYP) epileptiform spikes^24,27–30^. Prior in vitro studies reported that in seizures with LVF onset, activation of INs precedes that of PNs. In contrast to this view, in our data seizures with low fast activity (LVF) onset (34 of all 61 recorded seizures), we still observed the two initiation dynamics. 44.1% of seizures with LVF onset showed the INs-first initiation dynamics, while the remaining 55.9% showed simultaneous PNs-INs initiation dynamics. Comparing the duration of seizures revealed no significant differences between seizures with simultaneous PNs-INs and INs-first initiation dynamics (Fig. 2f).

Recruit synchrony at seizure onset was significantly higher in PV-expressing INs than PNs. However, as seizures progressed one second after seizure onset, inter-neuronal synchronization flipped (Fig. 2g). Furthermore, unlike PNs, where synchronization increased during the first second, synchronization between PV-expressing INs decreased during the initial second of the seizure (Fig. 2g). Onset recruitment synchrony of both PNs and INs was significantly higher in seizures with simultaneous PNs-INs initiation dynamics, however, these differences disappeared by 1 second after seizure onset (Fig. 2h,i).

We then investigated the relative activation of PNs and SST-expressing INs during the onset of 4-AP induced seizures (Fig. 3). Since the stratum pyramidale contains only few SST-expressing interneurons, we alternated calcium imaging between two imaging planes: the stratum pyramidale and the stratum oriens using an electrically tunable lens (ETL) at approximately 10Hz (Fig. 3a-c). Quasi-simultaneous calcium imaging recordings of CA1 PNs and SST-expressing INs again revealed the two distinct initiation dynamics. In 63.8% of seizures (30 seizures in 5 mice), SST-expressing INs and PNs were activated simultaneously (simultaneous PNs-INs initiation dynamics). Conversely, in the remaining 36.2% of seizures (17 seizures), activation of SST-expressing INs preceded the activation of PNs by an average of 0.89±1.06 seconds (INs-first initiation dynamics; Fig. 3a-d). Among the five recorded mice, two displayed a single simultaneous PNs-INs seizure initiation dynamics in all their recorded seizures, while the remaining three mice exhibited the two patterns alternately in different seizures (Fig. 3e). Importantly, similar to PV-expressing INs both initiation dynamics were also observed when examining only seizures with LVF onset^33–35^. Specifically, in 14 of 32 seizures with LVF onset (43.8%) INs-first initiation dynamics occurred, while the remaining 56.2% (18 seizures) showed simultaneous PNs-INs initiation dynamics.

**Figure 3.**
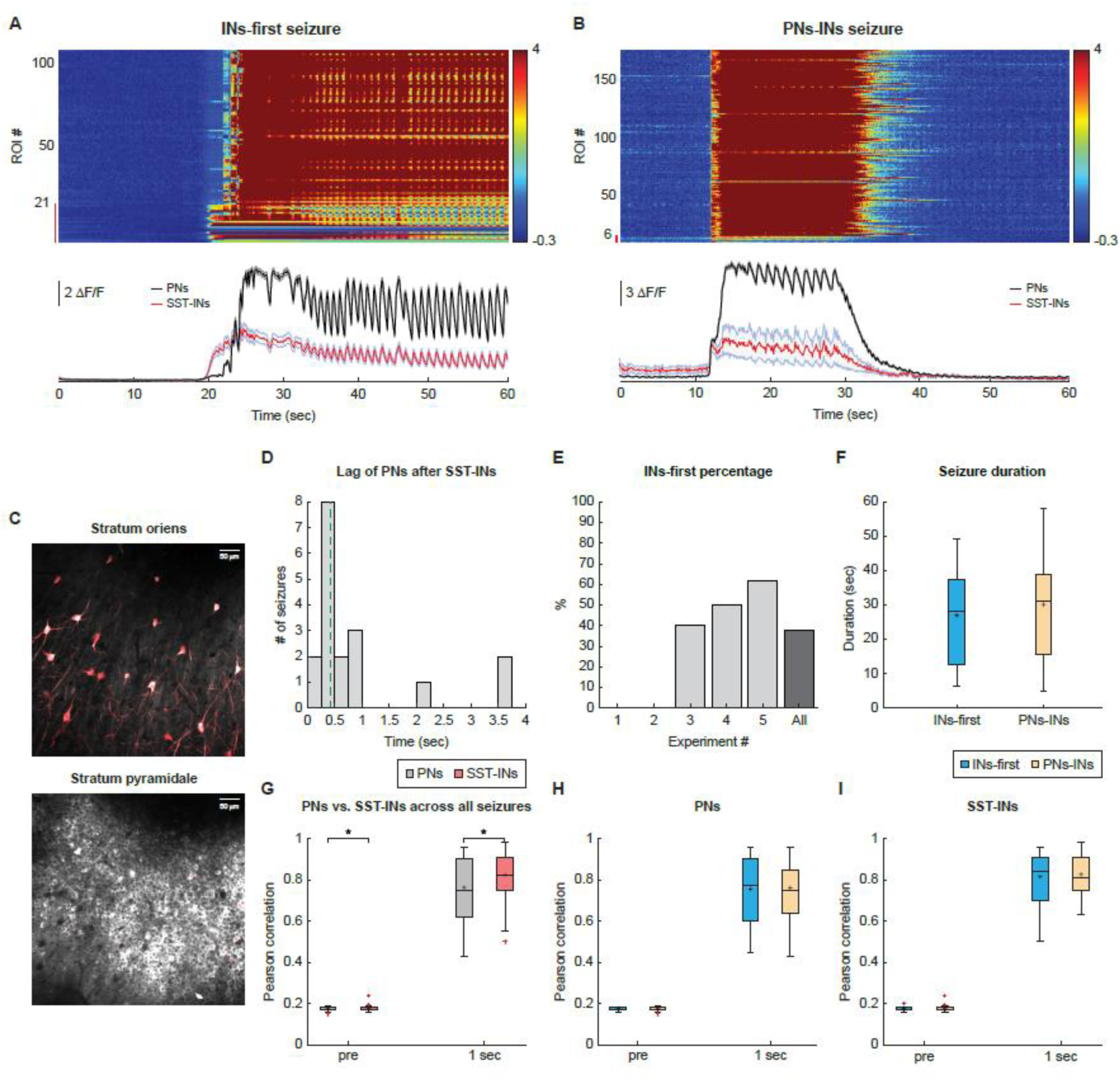
Simultaneous recordings of excitatory pyramidal neurons and inhibitory SST-expressing interneurons reveal dual dynamics during initiation of acute 4-AP induced seizures. **a,b,** The calcium fluorescence transients presented as ΔF/F of the individual SST-expressing INs (lower 21 ROIs marked by a red vertical line on the left) and PNs (all remaining ROIs) during two individual acute 4-AP induced seizures. The calcium transients were recorded using the calcium indicator GCamP6f and presented as color coded ΔF/F responses. The panels below show traces of the average (mean±SD) ΔF/F response over all recorded SST-expressing INs (red trace) and all simultaneously recorded PNs (black trace). Note that in the seizure in panel a, activation of SST-expressing INs preceded that of PNs (INs-first initiation dynamics), while in the seizure shown in panel b, PNs and SST-expressing INs were activated simultaneously (simultaneous PNs-INs initiation dynamics). c, Overlaid two-photon images of GCaMP6f which was expressed in both PNs and SST-expressing INs (gray), and of mRuby2-postive SST-expressing INs (red). Upper panel shows images from the stratum oriens where only IN somas are found, and lower panel shows images from the stratum pyramidale which is dominated by somas of PNs. d, A histogram showing the latency between initial firing of SST-expressing INs and PNs across all seizures with INs-first initiation dynamics. Dashed line marks the median value. e, A histogram of the percentage of seizures with INs-first initiation dynamics in the 5 different recorded mice (light gray) and average value for all seizures in all mice (dark gray right bar). f, Box plots of seizure duration of seizures with INs-first and simultaneous PNs-INs onset dynamics. g, Box plots of the average Pearson correlation coefficients of PNs compared to SST-expressing INs before and during the first second of seizures. h, Box plots comparing the average Pearson correlation coefficients of PNs before and during the first second of seizures with INs-first (blue) and simultaneous PNs-INs (yellow) initiation dynamics. i, Same as h for SST-expressing INs. PNs = pyramidal neurons, SST = Somatostatin, INs = interneurons, ROI = region of interest; *p < 0.05; transverse lines mark the median values, black crosses mark the mean values, and red crosses mark outliers.

Comparison of seizure duration revealed no significant differences between seizures with the two initiation dynamics (Fig. 3f). SST-expressing INs showed significantly higher synchronization compared to PNs one second after seizure onset (Fig. 3g), while no significant differences were seen in synchronization of either PNs or INs between the two seizure types (Fig. 3h,i). Unfortunately, the lower acquisition rate of the SST experiments did not allow us to reliably determine the inter-neural synchronization at seizure onset.

Previous studies have suggested that recruitment of PNs in seizures results from a depolarizing block of INs activity. Our experimental findings contradict this hypothesis.

Instead, we consistently observed sustained firing of both PV- and SST-expressing INs throughout seizures with INs-first initiation dynamics, persisting even as PNs were recruited (Fig. 2a,b, 3a,b).

### Relative activation of excitatory pyramidal neurons and inhibitory interneurons during initiation of spontaneous seizures in the chronic KA model

We conducted simultaneous calcium imaging recordings from both PNs and INs during the onset of spontaneous seizures in mice with the chronic KA model of epilepsy (Fig. 4). In contrast to acute 4-AP induced seizures, in the chronic KA model all recorded seizures showed simultaneous co-activation of PV-expressing INs and PNs (Fig. 4a, 34 seizures in 4 PV-Cre mice, 17 seizures in 2 SST-Cre mice).

**Figure 4.**
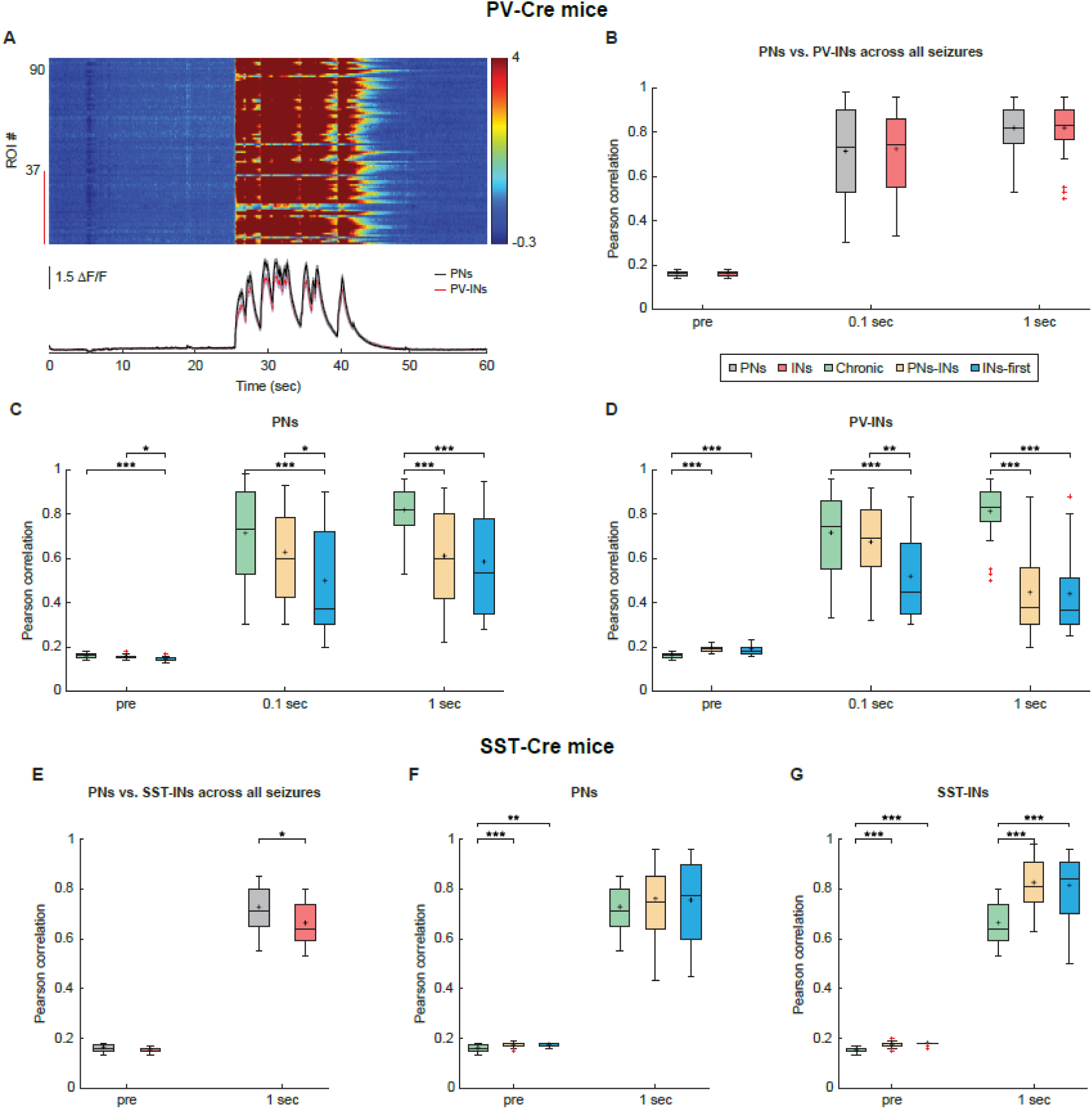
Simultaneous recordings of excitatory pyramidal neurons and inhibitory PV-expressing interneurons during spontaneous seizures in the chronic KA model of epilepsy. **a,** The calcium fluorescence transients presented as ΔF/F of the individual PV-expressing INs (lower 37 ROIs marked by a red vertical line on the left) and PNs (all remaining ROIs) during a spontaneous seizure from a mouse with KA induced chronic TLE. The calcium transients were recorded using the calcium indicator GCamP6f and presented as color coded ΔF/F responses. The panel below shows traces of the average (mean±SD) ΔF/F response over all recorded PV-expressing INs (red) and all simultaneously recorded PNs (black). Note that PNs and PV-expressing INs were activated simultaneously at the onset of the seizure. b, Box plots of the Pearson correlation coefficients of PNs (gray) and PV-expressing INs (red) during different time windows of seizures. c, Box plots of the Pearson correlation coefficients of PNs during different time windows of chronic spontaneous seizures (green), and acute 4-AP induced seizures with either INs-first (blue) or simultaneous PNs-INs (yellow) initiation dynamics. d, Same as c for PV-expressing INs. e, Box plots of the average Pearson correlation coefficients of PNs (gray) and SST-expressing INs (red) before and during the first second of spontaneous seizure in a mouse with the chronic KA model of epilepsy. f, Box plots of the average Pearson correlation coefficients of PNs before and during the first second of chronic spontaneous seizures (green), and acute 4-AP induced seizures with either INs-first (blue) or simultaneous PNs-INs (yellow) initiation dynamics. g, Same as f for SST-expressing INs. PNs = pyramidal neurons, PV = parvalbumin, SST = Somatostatin, INs = interneurons, ROI = region of interest; *p < 0.05, **p < 0.01, ***p < 0.001; transverse lines mark the median values, black crosses mark the mean values, and red crosses mark outliers.

Importantly, unlike 4-AP induced seizures where we likely recorded from the near vicinity of the seizure onset site in the chronic KA model the entire hippocampus was histologically affected, and the site of seizure onset was unclear. Thus, we may have recorded from propagated sites. This might explain why we only observed simultaneous PNs-INs initiation dynamics in the chronic KA model. A second possible explanation of the differences we observed between the acute and chronic models is the different epileptogenic process. While 4-AP increase firing of action potentials in axons^26–28^, chronic KA model seizures probably result from sprouting and increased connectivity between the surviving neurons^36,37^.

Synchronization of PNs and PV-expressing INs was not different at the onset of seizures (Fig. 4b). Moreover, synchronization of both PNs and PV-expressing INs was significantly higher in the chronic versus the acute model at both seizure onset. Interestingly, onset synchronization of chronic seizures more closely resembled acute seizures with simultaneous PNs-INs initiation dynamics (Fig. 4c,d).

Comparing activation of PNs and SST-expressing INs during onset of chronic spontaneous KA-induced seizures revealed synchronization was significantly higher in PNs in the first second of the seizure (Fig. 4e). Onset synchronization of SST-expressing INs in the chronic model was lower than that in acute 4-AP induced seizures with both initiation dynamics types (Fig. 4f,g).

### Chemogenetic inhibition of PV- and SST-expressing interneurons

We next aimed to investigate the causal relationship between INs activation and seizure onset by using chemogenetic and optogenetic manipulations. Suppressing the activity of INs by expressing inhibitory Gi (hM4Di) designer receptors exclusively activated by designer drugs (DREADDs)^38,39^ in either PV-expressing (Fig. 5a-d) or SST-expressing (Fig. 5e-h) INs had a significant pro-ictogenic effect. Suppression of both IN subtypes significantly shortened the latency from application of 4-AP to the first seizure and increased seizure frequency (Fig. 4b,c, 4f,g).

**Figure 5.**
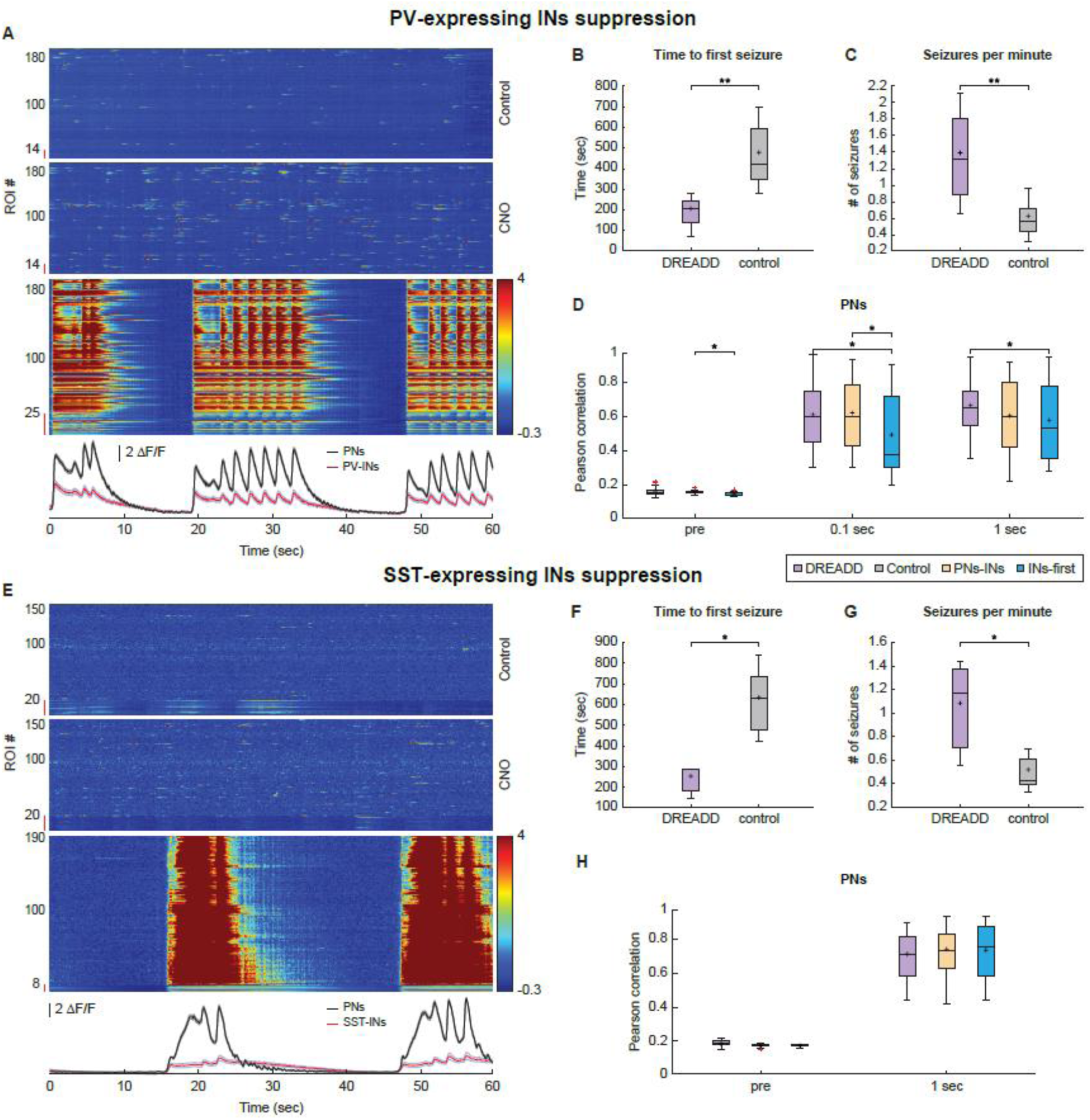
The ictogenic effect of chemogenetic silencing of PV- and SST-expressing interneurons. **a,** Three panels showing heat maps of the calcium activity from the same PV-expressing INs (ROIs adjacent to red vertical line) and PNs (all other ROIs) under control conditions (upper panel), after application of CNO to inhibit PV-expressing INs (middle panel), and during 4-AP induced seizures in the presence of CNO (lower panel). Color encodes the percent change in fluorescence (ΔF/F). The panel below shows the corresponding average (mean±SD) ΔF/F traces of all PNs (black) and all PV-expressing INs (red) during the seizures. Note that the activation of PNs and PV-expressing INs occurred simultaneously. **b,** Box plots of the time lag between 4-AP application and initiation of the first 4-AP induced seizure in control conditions and after DREADDs mediated inhibition of PV-expressing INs. **c,** Box plots of the frequency of 4-AP induced seizures in control conditions and after DREADDs mediated inhibition of PV-expressing INs. **d,** Box plots comparing the average Pearson correlation coefficients of PNs during 4-AP induced seizures under control conditions and in the presence of DREADDs mediated inhibition of PV-expressing INs. Seizures recorded in control condition are divided according to their initiation dynamics (INs-first or simultaneous PNs-INs). **e-f,** Same as **a-d** for DREADDs mediated inhibition of SST-expressing INs. PNs = pyramidal neurons, PV = parvalbumin, INs = interneurons, SST = somatostatin, ROI = region of interest; *p < 0.05, **p < 0.01; transverse lines mark the median values, black crosses mark the mean values, and red crosses mark outliers.

After DREADD-mediated suppression of PV-expressing INs all seizures exhibited simultaneous PNs-INs initiation dynamics (81 seizures in 4 PV-Cre mice and 61 seizures in 3 SST-Cre mice). When SST-expressing INs were suppressed, most seizures (42 out of 61 seizures, 68.9%) showed simultaneous PNs-INs initiation dynamics, while the remaining 19 seizures (31.1%) exhibited a new onset pattern of PNs-first dynamics, where firing of PNs preceded that of SST-expressing INs. Taken together the DREADDs experiments showed that early activation of INs is not a prerequisite for seizure initiation.

### Optogenetic activation of pyramidal neurons and PV- and SST-expressing interneurons

We next conducted optogenetic experiments wherein we explored the impact of selectively activating different cell types on seizure initiation. Specifically, we targeted three cell types: PV- and SST-expressing inhibitory INs, and CA1 excitatory PNs. To selectively activate either PV- or SST-expressing INs, we employed a Cre-dependent expression strategy of Channelrhodopsin-2 (ChR2) in PV- or SST-Cre transgenic mice. For selective activation of PNs, we utilized ChR2 expression under the control of the CaMK2 promoter. To evaluate the effect of optogenetic stimulation on ictogenesis, we analyzed seizure initiation across three different time segments: during the 10-second optogenetic stimulation, in the immediate post-stimulation period (5 seconds after the opto-stimulation ended), and during the unstimulated period (more than 20 seconds after stimulation terminated). Our findings revealed that optogenetic activation of both PV- (Fig. 6a-d) and SST-expressing INs (Fig. 6e-h) had pro-ictogenic effects. Activation of PV-expressing INs led to a 2.5-fold increase and activation of SST-expressing INs resulted in a 4-fold increase in the averaged probability of seizure initiation during opto-stimulation compared to the unstimulated period (Fig. 6c). Interestingly, only activation of PV-expressing INs and not SST-expressing INs increased seizure initiation in the immediate post-stimulation period (Fig. 6c,g), suggesting post-stimulation rebound activation occurs mostly after synchronized activation of PV-expressing INs and not SST-expressing INs. During both opto-stimulation of PV- and SST-expressing INs synchronization of PNs was not significantly different at the onset of seizures that initiated during opto-stimulation and the unstimulated period (Fig. 6d,h). Optogenetic activation of excitatory CA1 PNs also had pro-ictogenic effects and resulted in a 7.7-fold increase in seizure initiation during opto-stimulation as compared to the unstimulated period (Fig. 6i).

**Figure 6.**
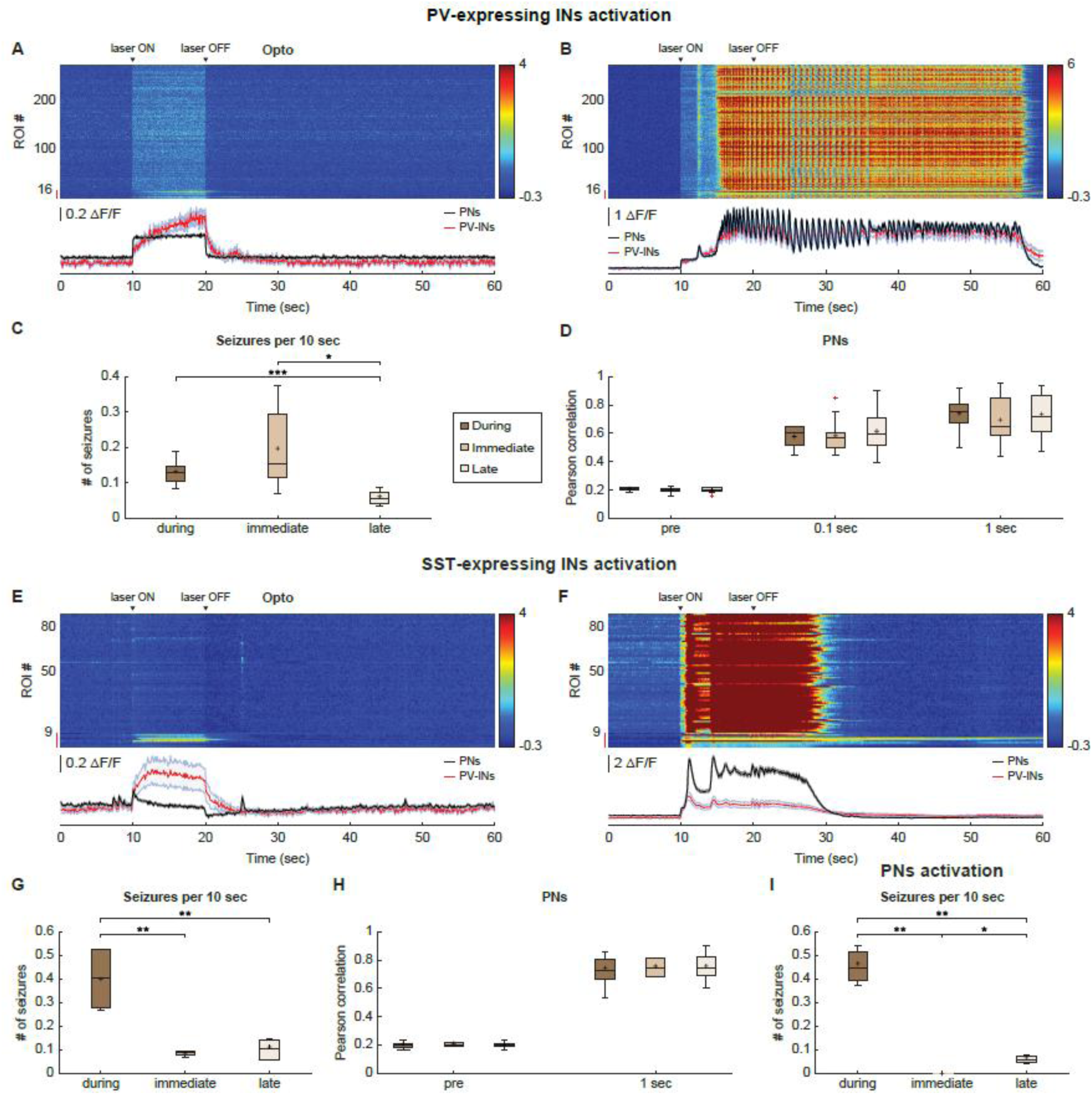
The ictogenic effect of optogenetic activation of excitatory pyramidal neurons, and PV and SST-expressing inhibitory interneurons. **a,b,** Heat maps of the calcium activity recorded from PV-expressing INs (ROIs adjacent to red vertical line) and PNs (all other ROIs) before, during and after a 10 second optogenetic activation of PV-expressing INs. Color in the heat map encodes the percent change in fluorescence (ΔF/F). The panels below show the corresponding traces of the average (mean±SD) ΔF/F of all PNs (black) and of PV-expressing INs (red). Panel a shows a trace with no seizure. Note the gradual increase of activation in PV-expressing INs during optogenetic stimulation. Panel b shows a trace with a seizure evoked during the optogenetic activation of PV-expressing INs. c, Box plots of seizure frequency (per 10 seconds) occurring during three different time windows of the trace: during the optogenetic stimulation of PV-expressing INs, immediately after stimulation ended (first 5 seconds post-stimulation) and late after the cessation of optogenetic stimulation (at least 20 seconds after the stimulation ended). d, Box plots comparing the average Pearson correlation coefficients of PNs during the initial 100 milliseconds of seizures evoked during optogenetic stimulation of PV-expressing INs, immediately after the stimulation ended (first 5 seconds post-stimulation) and late after cessation of optogenetic stimulation (at least 20 seconds after the stimulation ended). e-h, Same as a-d for optogenetic activation of SST-expressing INs. i, Same as c for the optogenetic activation of PNs. PNs = pyramidal neurons, PV = parvalbumin, INs = interneurons, SST = somatostatin, ROI = region of interest; *p < 0.05, **p < 0.01, ***p < 0.001; transverse lines mark the median values, black crosses mark the mean values, and red crosses mark outliers.

In summary, our chemogenetic and optogenetic experiments support the notion that several distinct dynamics can lead to initiation of temporal lobe seizures. Specifically, both activation and inhibition of PV-expressing or SST-expressing INs and activation of PNs can each independently lead to seizure initiation.

### Recruitment Robustness of pyramidal neurons and interneurons during onset of seizures

Next, we investigated whether seizure initiation follows a robust and stereotypic route. To address this question, we determined the recruitment rank order of PNs and INs at the onset of each individual seizure. We then assessed the consistency of the recruitment patterns during seizure onset by calculating a matrix of the pairwise Spearman correlation coefficients between all possible pairs of seizures across all different seizures in the same mouse and then obtained the average value for each mouse (Fig. 7 and 8). Neuronal activation was defined as a calcium response amplitude exceeding three standard deviations above the pre-seizure baseline.

**Figure 7.**
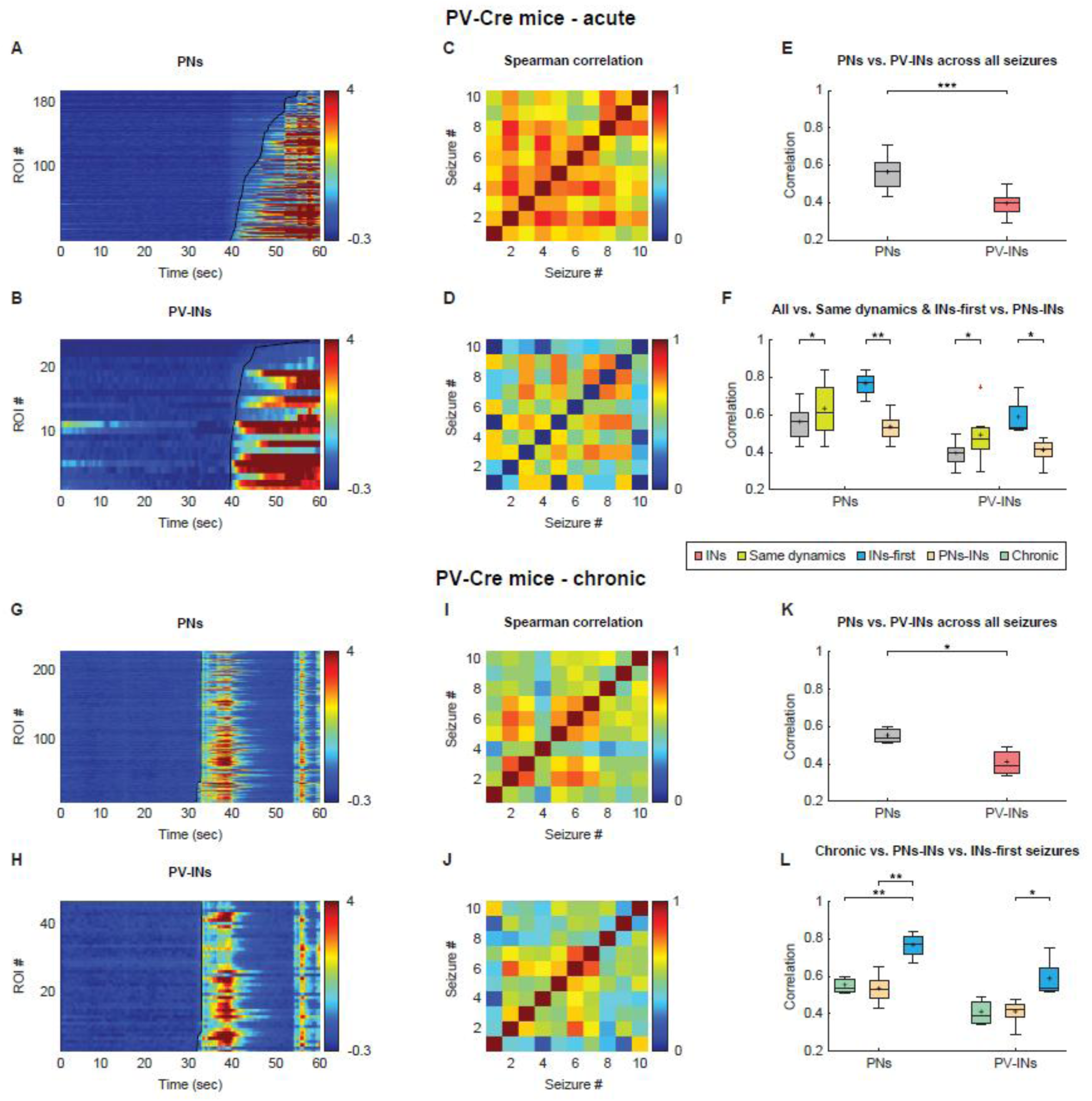
Recruitment of excitatory pyramidal neurons and PV-expressing inhibitory interneurons at the onset of seizures in acute and chronic epilepsy. **a**, The calcium fluorescence transients presented as ΔF/F for the individual PNs during a 4-AP induced seizure. The calcium transients were recorded using the calcium indicator GCamP6f, presented as color coded ΔF/F responses, and arranged by the temporal activation order of the ROIs (neurons). Black line marks the onset time of activation of the different ROIs. b, Same as a for the PV-expressing INs during the same seizure shown in a. c, The inter-seizure Spearman correlation matrix for recruitment of PNs at the onset of 4-AP induced seizures. Each box in the matrix represents the Spearman correlation coefficient between the recruitment order of PNs of a pair of seizures, and the matrix represents the correlations between all seizure pairs in the same mouse (in this case 10 different seizures). d, Same as c for recruitment of PV-expressing INs in the same mouse. e, Box plots of the average inter-seizure Spearman correlation coefficients of the temporal recruitment of individual neurons averaged initially for all 4-AP induced seizures in each mouse and then further averaged over all mice for PNs (gray) and PV-expressing INs (red). f. Box plots comparing the average inter-seizure Spearman correlation coefficients of the temporal recruitment of individual neurons in seizures in four conditions: all seizures (gray), seizures with similar initiation dynamics (either INs-first or simultaneous PNs-INs (green)), only seizures with INs-first initiation dynamics (blue), and only seizures with simultaneous PNs-INs initiation dynamics (yellow). The data is shown for PNs (left) and PV-expressing INs (right). The Spearman correlation coefficients were initially averaged for seizures in individual mice and then further averaged over all mice. g-l, Same as a-f for spontaneous seizures in mice with the chronic KA epilepsy model. Note that in l, data from chronic spontaneous seizures (green) is compared to acute 4-AP induced seizures with either INs-first (blue) or simultaneous PNs-INs initiation dynamics (yellow). PNs = pyramidal neurons, PV = parvalbumin, INs = interneurons, ROI = region of interest; *p < 0.05, **p < 0.01, ***p < 0.001; transverse lines mark the median values, black crosses mark the mean values.

**Figure 8.**
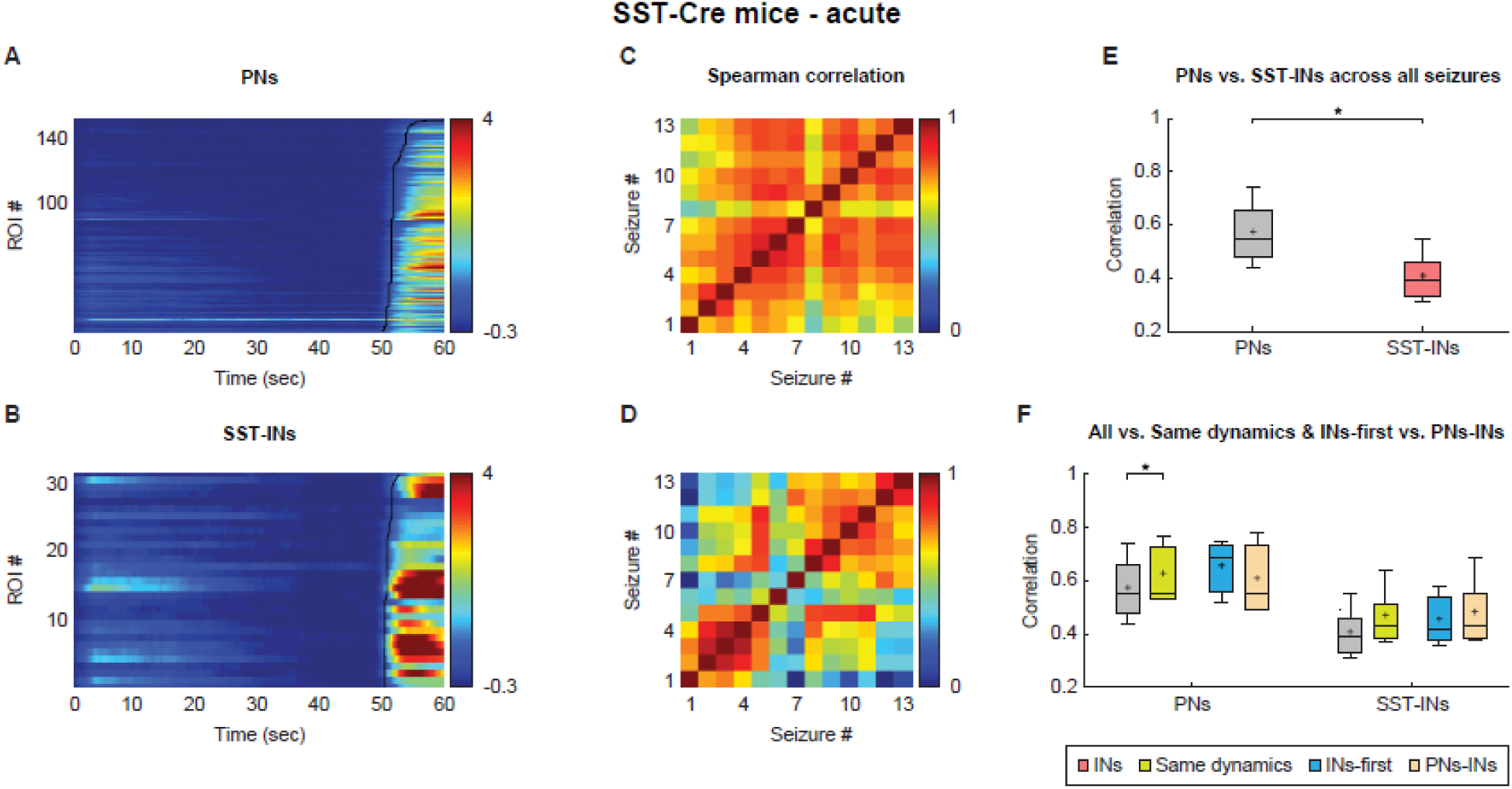
Recruitment of pyramidal neurons and SST-expressing interneurons at the onset of 4-AP induced seizures in acute epilepsy. **a**, The calcium fluorescence transients presented as ΔF/F for the individual PNs during a 4-AP induced seizure. The calcium transients were recorded using the calcium indicator GCamP6f, presented as color coded ΔF/F responses, and arranged by the temporal activation order of the ROIs (neurons). Black line marks the onset time of activation of the different ROIs. b, Same as a for the SST-expressing INs during the same seizure shown in a. c, The inter-seizure Spearman correlation matrix for recruitment of PNs at the onset of 4-AP induced seizures. Each box in the matrix represents the Spearman correlation coefficient between the recruitment order of PNs of a pair of seizures, and the matrix represents the correlations between all seizure pairs (in this case 13 different seizures) in the same mouse. d, Same as c for SST-expressing INs in the same mouse. e, Box plots of the average inter-seizure Spearman correlation coefficient of the temporal recruitment of individual neurons averaged initially for all 4-AP induced seizures in each mouse and then further averaged over all mice for PNs (gray) and SST-expressing INs (red). f, Box plots comparing the average inter-seizure Spearman correlation coefficients of the temporal recruitment of individual neurons in seizures in four conditions: all seizures (gray), seizures with similar initiation dynamics (either INs-first or simultaneous PNs-INs (green)), only seizures with INs-first initiation dynamics (blue), and only seizures with simultaneous PNs-INs initiation dynamics (yellow). The data is shown for PNs (left) and SST-expressing INs (right). The Spearman correlation values were initially averaged for seizures in individual mice and then further averaged over all mice. PNs = pyramidal neurons, SST = somatostatin, INs = interneurons, ROI = region of interest; *p < 0.05; transverse lines mark the median values, black crosses mark the mean values.

We first examined recruitment of PNs and PV-expressing INs during acute and chronic seizures (Fig. 7). We observed significant similarities in the recruitment order of PNs across different seizures. This held true for both 4-AP-induced seizures (Fig. 7a-f) and spontaneous seizures in the chronic KA model (Fig. 7g-l). The average Spearman correlation coefficient of PNs recruitment in different seizures was 0.56±0.1 for acute 4-AP-induced seizures (8 mice) and 0.52±0.1 for spontaneous seizures in the chronic KA model (4 mice). Thus, the recruitment of CA1 PNs during seizure onset tended to replicate across different seizures in both the acute 4-AP and chronic KA models. These findings suggest that PN recruitment during seizure onset follows an orderly predictable path and is driven by the activation of the same group of neurons in a relatively robust and orderly manner. The recruitment order of PV-expressing INs showed less consistency across seizures compared to PNs in both acute 4-AP-induced seizures (Fig. 7e; average Spearman correlation coefficient of 0.39 ± 0.06 in 8 mice) and spontaneous seizures in the chronic KA model (Fig. 7k; average Spearman correlation coefficient: 0.41 ± 0.07 in 4 mice). When comparing the recruitment robustness of seizures with the two different initiation dynamics we found higher Spearman correlation coefficients in both PNs and INs of seizures with INs-first initiation dynamics. Moreover, the consistency of the recruitment order of PNs and PV-expressing INs significantly increased in seizures that exhibited the same initiation dynamics, either INs-first or simultaneous PNs-INs initiation patterns, as compared to the case where all seizures were averaged regardless of their initiation dynamics (Fig. 7f,l). These findings further highlight key differences in initiation mechanisms between seizures with distinct initiation profiles.

Similar results were also observed for acute 4-AP induced seizures for SST-expressing INs (Fig. 8). Due to the limited data available for SST-expressing INs in the chronic KA model (2 mice), we were unable to derive average results of SST-expressing INs recruitment during chronic spontaneous seizures.

## Discussion

In this study, we investigated the mechanisms underlying initiation of focal temporal lobe seizures in vivo using two different models of TLE: acute 4-AP induced seizures and spontaneous seizures in the chronic KA model. To our knowledge, this study represents the first exploration of ictogenesis in the hippocampus in vivo with such fine-grained resolution.

The key findings of this study include: (1) We identified two different network dynamics that lead to seizure initiation. The first involves initial activation of inhibitory INs followed by the recruitment of excitatory PNs. The second entails rapid activation of both PNs and INs, which likely reflects the initial activation of excitatory PNs followed by rapid recruitment of inhibitory INs due to rapid feedback mechanisms. Interestingly both dynamics occur in different seizures generated within the same network of the same mouse. This conclusion is supported by both our simultaneous recordings of PNs and INs during seizure initiation, and by our opto-and chemogenetic experiments demonstrating that both activation and suppression of INs, as well as the activation of PNs, enhance seizure initiation. (2) We assessed the robustness of seizure initiation at the single neuron level, and found the recruitment rank order of neurons during seizure onset to be robust between different seizures. This was particularly evident for PNs. These findings indicate that the initiation of seizures is an orderly and stereotypic process, characterized by a relatively robust recruitment pathway of neurons within the epileptogenic network.

While ictogenesis of temporal lobe seizures has been extensively studied in the past, almost all these studies have been conducted in vitro, particularly in brain slices^40–42^. Fewer studies have been performed on isolated whole-brain in vitro preparations^43,44^, and only a handful of studies have explored the E/I interactions during the initiation of TLE. These studies utilized single unit recordings^45,46^, which is a less precise method than two-photon calcium imaging, and cannot differentiate between subtypes of INs.

Our findings reveal that 30-40% of 4-AP induced seizures exhibit an initial increase in firing of inhibitory INs, followed by a subsequent activation of excitatory PNs (INs-first initiation dynamics). These findings align with prior observations reported in 4-AP induced seizures in hippocampal and hippocampal-entorhinal cortex brain slices, as well as in isolated brain preparations in vitro^3–6,40–42^. However, in contrast to these studies we found that INs-first dynamics occurs only in a minority of acute 4-AP induced seizures, while the majority of seizures showed simultaneous activation of PNs and INs. Importantly this also held true for seizures with LVF onset.

Our calcium imaging recordings were acquired at 30Hz, limiting our ability to detect time differences smaller than 50-100 milliseconds, Therefore, seizures with simultaneous PNs-INs initiation dynamics should be interpreted within this limitation. It is probable that in fact PNs were activated first, followed by a rapid secondary activation of INs due to robust feedback connections between PNs and INs, albeit with a time difference smaller than our detection threshold.

Importantly, simultaneous PNs-INs initiation dynamics were also observed in seizures with LVF onset, and not only in seizures showing a HYP onset pattern^32–35^. Interestingly, previous in vitro studies did report some seizures demonstrating initial activation of PNs^8,45–47^, but the onset pattern of these seizures exhibited HYP and not LVF onset.

There are several prevailing hypotheses regarding the mechanisms underlying the unexpected discovery of inhibitory INs driving seizure initiation. These include the possibility that intense firing of INs leads to accumulation of extracellular potassium which increases excitability of PNs and/or accumulation of intracellular chloride which reduces the inhibitory efficacy of GABA-A receptors. Other possibilities include depolarizing block of IN firing and/or synchronized rebound activation of PNs following the early intense activation of INs^3,4,13–15,17,48–50^. Our experiments failed to observe either phenomena.

The differences between our findings showing dual initiation dynamics and previous studies in vitro, which primarily found INs-first onset initiation, can be attributed to the disparity between in vivo and in vitro preparations, or the possibility that we may be recording from propagated sites and not the site of seizure onset. It is important to note that prior studies using 4-AP induced seizures in combined entorhinal-hippocampal brain slices revealed seizures initiate in the entorhinal cortex^3,40^, while in the slice culture preparation seizures initiated in the dentate gyrus^16^. In the case of recordings from propagated sites INs-first dynamics may signify an inhibitory restraint developing in response to ictal activation of PNs in adjacent areas^51,52^.

In contrast to our findings in the chronic KA model, previous studies in the chronic pilocarpine model in vivo reported firing of INs preceded that of PNs^45,46^ at seizure onset. These differences probably reflect the different recording methods, two-photon calcium imaging versus single unit recordings, and the different methods for cell type identification. To the best of our knowledge our study is the first to examine ictogenesis in a chronic TLE model using two-photon calcium imaging in vivo.

In this study, we compared activation of excitatory PNs and two types of inhibitory INs: PV-expressing INs and SST-expressing INs, which primarily target the axo-somatic region and the dendrites, respectively^30,31^. Our recordings revealed dual initiation dynamics, with both subtypes demonstrating a comparable incidence of seizures (30-40%) exhibiting INs-first initiation dynamics. It is worth noting, however, that PV-expressing INs preceded PNs activation by a longer time lag compared to SST-expressing INs. These findings suggest that seizures characterized by INs-first initiation dynamics initially involve activation of PV-expressing INs, followed by activation of SST-expressing INs, and ultimately the activation of PNs.

Comparison of our findings in the CA1 region with previous studies conducted in the neocortex, using a similar epilepsy model and recording techniques, reveals significant differences. Unlike acute 4-AP induced temporal lobe seizures which demonstrated dual dynamics of ictogenesis, 4-AP induced neocortical seizures exhibited only a single initiation dynamics of simultaneous PNs-INs initiation dynamics in layer 2/3^17^. This difference may reflect dissimilarities in the local network structure and cell density between the neocortex and hippocampus.

One of the major challenges we face is to go beyond the observational phase and define causality^10,11^. To address this question, we used opto-and chemogenetic perturbations, which further supported the possibility that two different routes can lead to seizure initiation. We found that optogenetic activation of both PV- and SST-expressing INs and PNs all had pro-ictogenic effects. These findings are consistent with previous findings in temporal and neocortical epilepsy in vivo and in vitro^5–15,53–56^.

The second major question we deal with in this study is how stereotypical and robust is the ictogenic process. We found a high degree of correlation between the recruitment rank order of neurons during initiation of different seizures in the same mouse. These findings indicate that different seizures initiate stereotypically in an orderly manner within the same subgroup of neurons. This was true for both acute 4-AP induced seizures and spontaneous seizures in the chronic KA model. Thus, there is a small subgroup of neurons where seizures initiate, and from that initial group of “epileptic neurons” the seizure activity recruits additional neurons in a typical and orderly manner until the entire local network is involved. Interestingly, the inter-seizure robustness of the recruitment rank order was greater for PNs than INs. This may be related to the gap junctions that connect INs^56^. Moreover, the resemblance of the recruitment rank order of different seizures was greater for seizures with INs-first initiation dynamics, and for seizures that shared the same initiation dynamics, either INs-first or simultaneous PNs-INs.

Finally, it is important to stress our study concentrated on the network dynamics of ictogenesis in mesial TLE. Yet we did not address the cellular mechanisms nor the functional connectivity underlying epileptogenesis. The etiology of epileptogenesis and the location of the epileptogenic zone can influence the specific dynamics underlying ictogenesis, as demonstrated by the differences we found between mesial temporal and neocortical epilepsy. Further studies are required to tackle these important questions.

## Acknowledgment

This work was supported by the Israel Science Foundation (ISF), and the Prince Center for neurodegenerative disorders.

## Author Contributions

AM and YS contributed to the conception and design of the study; AM contributed to acquisition of the data; SA and YS contributed to analysis of data; and AM and YS contributed to drafting the text or preparing the figures.

